# Estimating RNA dynamics using one time point for one sample in a single-pulse metabolic labeling experiment

**DOI:** 10.1101/2020.05.01.071779

**Authors:** Micha Hersch, Adriano Biasini, Ana C. Marques, Sven Bergmann

## Abstract

Over the past decade, experimental procedures such as metabolic labeling for determining RNA turnover rates at the transcriptome-wide scale have been widely adopted and are now turning to single cell measurements. Several computational methods to estimate RNA processing and degradation rates from such experiments have been suggested, but they all require several RNA sequencing samples. Here we present a method that can estimate RNA synthesis, processing and degradation rates from a single sample. Our method is computationally efficient and outputs rates that correlate well with previously published data sets. Using it on a single sample, we were able to reproduce the observation that dynamic biological processes tend to involve genes with higher metabolic rates, while stable processes involve genes with lower rates. This supports the hypothesis that cells control not only the mRNA steady-state abundance, but also its responsiveness, i.e., how fast steady-state is reached. In addition to saving experimental work and computational time, having a sample-based rate estimation has several advantages. It does not require an error-prone normalization across samples and enables the use of replicates to estimate uncertainty and perform quality control. Finally the method and theoretical results described here are general enough to be useful in other contexts such as nucleotide conversion methods and single cell metabolic labeling experiments.

## 1 Introduction

Since the advent of molecular biology, a consensus has emerged that the regulation of gene expression underlies most biological processes including development, disease and adaptation [1, 2, 3]. While gene expression regulation has mostly been associated with activating the production of RNA (e.g. through transcription factors), it has become apparent that the regulation of RNA splicing and RNA stability also plays an important role in determining the expression level of a gene [4, 5]. Taking advantage of high throughput RNA quantification protocols, methods designed to distinguish the effects of RNA synthesis, processing and degradation at the transcriptome-wide level have been developed. Among them, RNA metabolic labeling techniques relying on chemically modified ribonucleotides such as 4-thiouridine (4sU) and 5’-Bromouridine (BrU) have been widely adopted, as their impact on cellular function is minimal [6]. Briefly, incubating cells with modified ribonucleotides for a limited period of time (referred to as the pulse), and their concomitant incorporation in newly synthesized transcripts, allows distinguishing newly transcribed from preexisting RNA, which can be biochemically separated and quantified. This quantification, initially performed using microarray [7] and later using RNA-seq [8, 9], can then be used to estimate RNA decay rates. More recently, methods that rely on nucleotide conversion have been used to the same effect, with the advantage of circumventing the cumbersome biochemical enrichment and separation step.

In the last decade, several methods to estimates RNA dynamics from metabolic labeling experiment data have been developed [10, 11, 12] (see [13] for a review). Typically, labeled transcript abundances are fitted to an exponential function approaching to steady-state equilibrium (during or after the pulse), from which the RNA half-life can be estimated [14, 15, 16]. This requires time-course experiments in order to have enough points for fitting, as well as a way to normalize RNA concentrations across samples, either using spike-ins [17], or using internal controls such as intron concentrations [18]. The INSPEcT method [19] goes beyond first order dynamics and takes into account the RNA processing rates, which are estimated along with the degradation and synthesis rates. This method works by first estimating rates for individual samples by assuming no degradation during the pulse and then uses those estimates as a starting point for fitting model of rate evolution for all the rates of all samples. Those methods rely, for each sample, on a the separate quantification of labeled RNA on one hand and of total (mixed labeled and unlabeled) and/or unlabeled (or preexisting) RNA on the other hand. In its later version, INSPEcT was extended to estimate rates without labeling the sample [20].

In this work, we build on the INSPEcT approach and derive an exact solution (when it exists) for the initial rate estimates without making the assumption of no labeled transcript degradation. This is achieved by considering the intron to exon ratio for each transcript in both the labeled and unlabeled RNA pools, thus allowing to bypass the need for normalization across those two samples. We can thus infer synthesis, processing and degradation rates from a single sample and time point. Those rates can be used as such, allowing to reduce the experimental load and costs and compare rates across samples and time points. But they can also be used, as in INSPEcT, as initial estimates for mutliple samplebased rate estimation. Applying our method to our own experimental data and using a single sample and time point, we obtain synthesis and processing rates that are well correlated with the ones obtained using INSPEcT first guess. The degradation rates, on the other hand, correlate poorly across the two methods, but those computed with our method correlate better than INSPEcT with previously published mRNA degradation rates obtained with three replicates and seven time points [21]. Because it can be reduced to numerically solving an equation with a single unknown on a bounded domain, it is also faster than INSPEcT. Moreover, our results are consistent with an adapted gene-specific mRNA responsiveness and co-transcriptional mRNA processing [22].

## 2 Method

### 2.1 Oveview

This paragraph summarizes the general strategy of the method, with references to relevant equations indicated in parentheses. We use the Zeisel model of RNA dynamics [23] to model both the unlabeled and the labeled RNA (1, 2). Using the standard procedure for solving systems of linear differential equations, we find its general solution and its free parameters by setting the initial conditions for both the unlabeled (or pre-existing) and the labeled RNA (3–6), as illustrated in Fig 1. We can then express, for a given gene, the ratios for both unlabeled and labeled RNA of intron to exon expression level as functions of the processing and degradation rate of that gene (8,9). These two ratios are independent from the RNA synthesis rate. Using the intron to exon ratios as observables, we are left with two non-linear equations and two unknowns, namely the processing and degradation rates. These equations are then reparametrised with dimensionless parameters and reduced to a single non-linear equation with one unknown (12). This resulting equation is only defined on a bounded domain (13). Our rates can thus be inferred by numerically solving that equation on a bounded domain, which is very fast. In addition, we prove in Appendix C that this equation, under certain conditions, has a single solution (but in general it can also have two or no solution).

**Figure 1:**
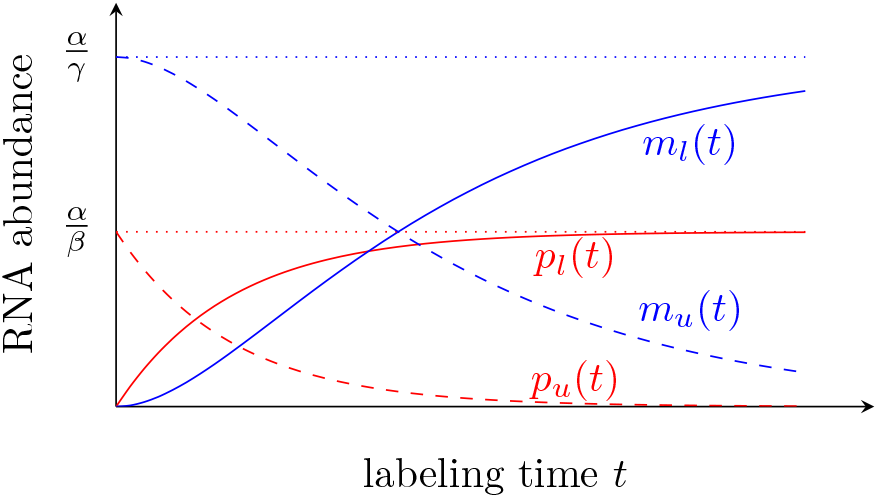
Evolution of unlabeled and labeled, premature and mature RNA during labeling according to the Zeisel model. Dotted horizontal lines correspond to steady-state levels, dashed lines correspond the unlabeled RNA and solid lines to labeled RNA. Processing and degradation rates can be estimated from the ratios of the two dashed lines and of the two solid lines at a single time point.

### 2.2 Model

Like previous work [19], we use the Zeisel model of RNA synthesis, processing and degradation [23].

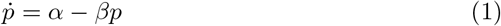

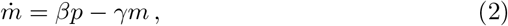

where *p* is the premature RNA, *m* the mature RNA, and *α*, *β, γ* are RNA the synthesis, processing and degradation rates. This model can be solved analytically (see appendix A). In particular, enforcing the boundary conditions corresponding to the unlabeled RNA, namely that it is at steady-state when the pulse starts (*t* = 0) and that subsequently no more pre-mature RNA is produced, results in

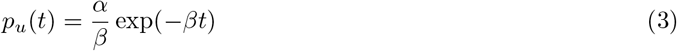

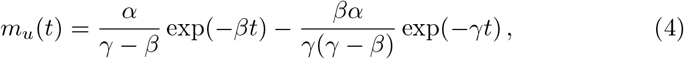

where the *u* subscript indicates that this corresponds to the unlabeled RNA pool.

Enforcing boundary conditions corresponding to the labeled RNA, namely that it is not (yet) expressed at *t* = 0 leads to

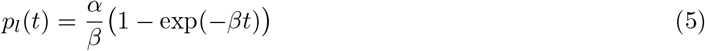

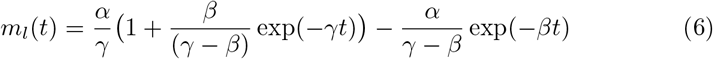

where the *l* subscript indicates that this corresponds to the labeled RNA pool.

### 2.3 Inferring synthesis, processing and degradation rates

We consider that the exonic RNA abundance *χ* corresponds to the premature and mature RNA, while the intronic RNA abundance *ι* correspond to the premature RNA only. Furthermore, we assume that χ and *ι* are suitably normalised for exonic and intronic length so that they are proportional to the number of transcripts. We can then compute:

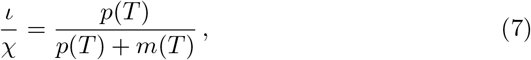

where *T* is the duration of the labeling.

In the case of unlabeled fraction, we have

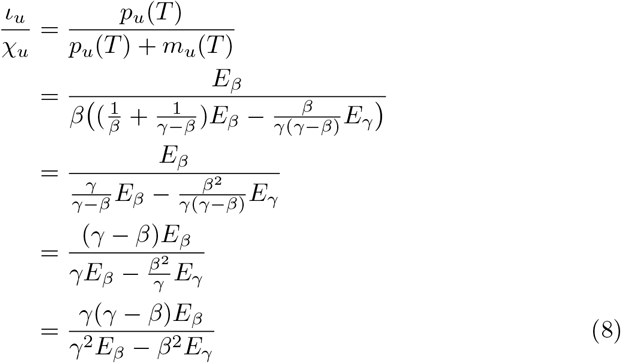

where we define *E_β_* = exp(−*βT*) and *E_γ_* = exp(−*γT*) as abbreviations.

For the labeled fraction, we have

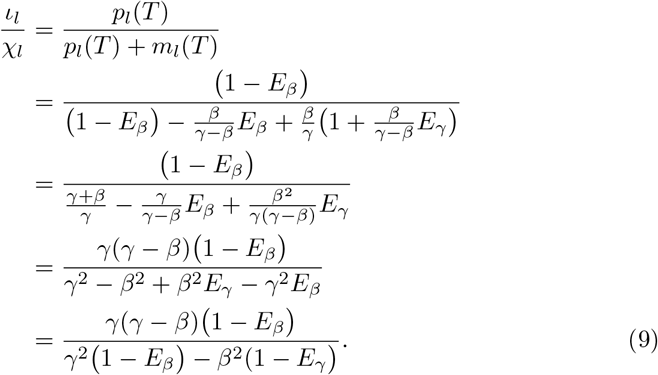

We notice that this last expression is of the same form as the one for the unlabeled fraction (8), but replacing exponentials by their complement to one. Importantly these two fractions do not depend on *α*, which (unlike [20]) allows our method to estimate processing and degradation rates independently from the synthesis rate.

Denoting 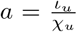 and 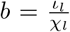 as the observable unlabeled and labeled fractions of intron abundance, we are left with a system of two equations and two unkowns *β* and *γ*, which we now set out to solve. First, we reparametrise our system with *β* = *kγ* and define *E_*kγ*_* = *E_*β*_* = exp(−*kγT*) leading to

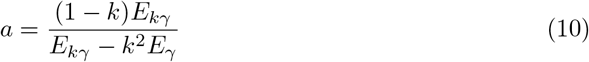

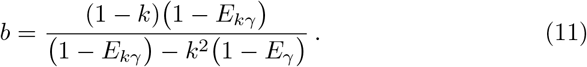

It is shown in Appendix B that system of equations can be simplified to the following equation in *k*:

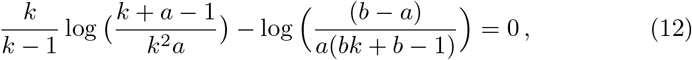

with the following domain of definition 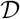 for *k*:

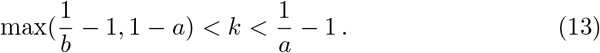

The above equation does not explicitely depend on *T* can be solved numerically on 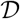. In practice *a* and *b* are approximated by *r_u_* and *r_l_*, defined as the length-normalized intronic to exonic read count ratio (or TPM ratio) for the unlabeled and for the labeled sampled respectively.

We further prove in Appendix C that for 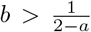, Eq. (12) has a single solution in the domain given by (13), which can be found very efficiently. This enables the estimation of the processing and degradation rates for a single sample. Moreover, since the reduced equation is independent from *T*, uncertainty on its true value does not affect the relative values of the resulting rates. Hence replicates can be used to assess the reliability of the estimates and time courses allow to test whether the rates are constant as assumed by the model.

If (12) does not have a solution, estimates can be obtained by minimizing (in log space) the squared Euclidean distance between the observed (i.e., *r_u_*, *r_l_*) and derived values of *a* and *b*:

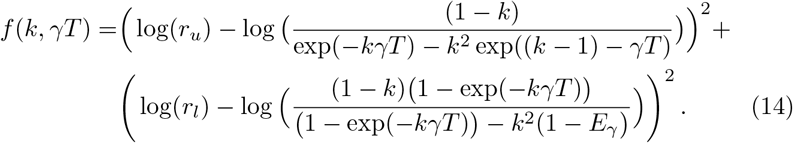

The ratios *r_u_*, *r_l_* must be smaller than one to make sense within our model and genes where this is not the case should be discarded. The log function is used to give exon and intron counts equal standing.

The above bivariate function can be reduced to a univariate function *f*\* using (36):

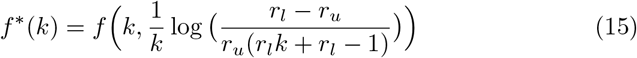

The processing and degradation rates are derived from *k* using (36) where *a* and *b* are again approximated by *r_u_* and *r_l_* respectively. Then the synthesis rate *α* can be easily obtained from (4), where *m_u_* is approximated by *χ_u_* (which is likely the most reliably measured specie):

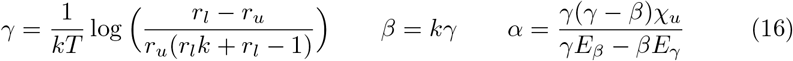

## 3 Results

### 3.1 Simulated data

In order to confirm that our method can be applied in principle, we evaluated our method on simulated data, where the data was generated using the exact model used to develop the method (see equations 3 and following). We chose not to simulate noise or biases in the data, as the aim of the simulation is only to validate the mathematical developments above and our implementation of the method. We generated 50000 random value for *α, β*, and *γ* ranging between exp(−5) and exp(5) and computed the corresponding values for *ι* and *χ*. We then computed *r_u_* and *r_l_* by taking the ratio. Estimates 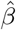 and 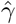 where then inferred by using *r_u_* and *r_ι_* as an input to the method and compare the original *β* and *γ*.

Numerically solving equation (12), yielded either one or two solutions. The results for the unambiguous cases are shown in Fig 2, left. We see that in virtually all cases, the method yields accurate estimates of the processing and degradation rates. For a few points, the method is less accurate at the upper boundary of the parameter space, probably due limited floating point precision. Indeed if the labeling time is too long with respect to the metabolic rates, virtually all unlabeled RNA are degraded and the rates cannot be reliably estimated.

**Figure 2:**
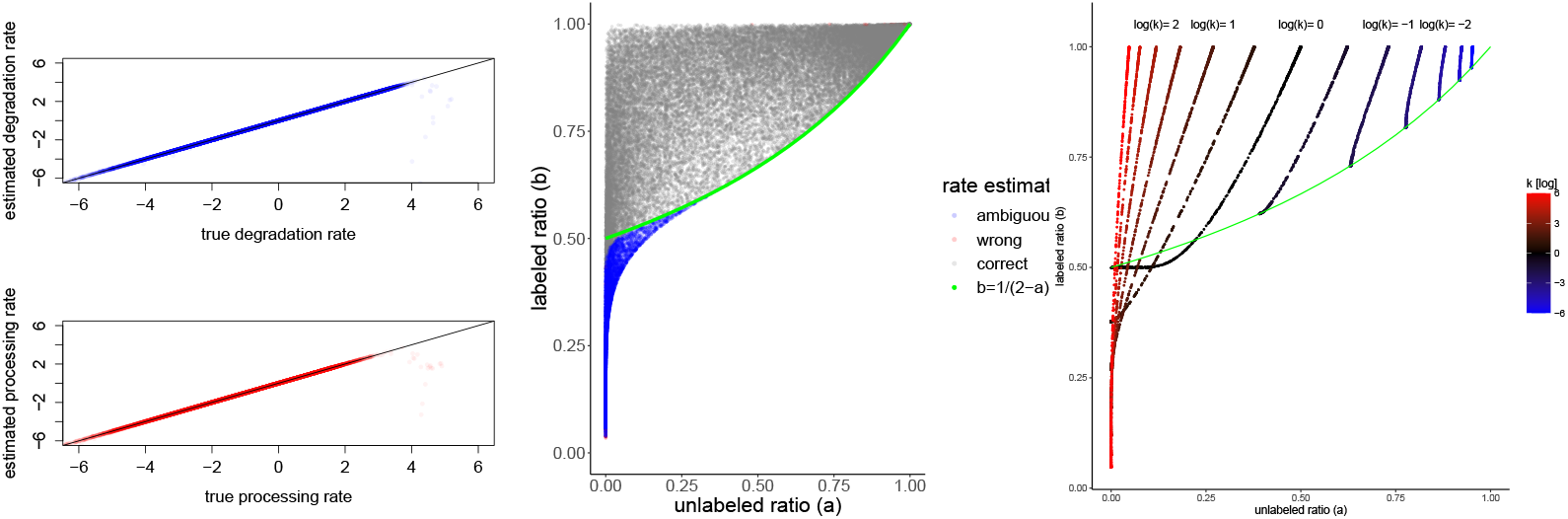
Simulated data. Left: the method correctly estimates processing and degradation rates. Points with ambiguous solutions are not shown. Some points corresponding to high rates cannot be estimated correctly as the system as already reached steady-state during the simulated “pulse”. Center: the measurement space can be partitioned into ambiguous and unambiguous regions. The green line corresponds to 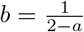. Above that line, rates are correctly and unambiguously estimated. Boundary cases are sometimes wrongly estimated, probably due to numerical errors (red dots). Right: Trajectories in the phase space are solely determined by the *k* parameter. They start at time *T* = 0 at the top (*b* =1) and go down. For *k* < 1 the trajectories (in blue) remain above the green line defined by *b* = (2 – *a*)^−1^ and do not cross. For *k* > 1 (in red), they cross each other below the system follows the trajectory depends on the actual values of *β* and *γ*.

As we are considering single-sample estimates, it is possible to chart the observable space given by *a* and *b* and see when the method provides unambiguous results. Fig. 2, center, confirms that for 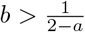 the method provides a unique (and correct) solution as proven in appendix C. Below this line (displayed in green), the methods provides ambiguous results as two distinct set of values *β* and *γ* can account for the same value of *a* and *b* (in blue). It is also possible to visualize the trajectories of the observables *a* and *b* for various values of *k*, as depicted in Fig. 2, right. When *T* = 0, trajectories start from the top of the space at 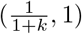. When *k* < 1, as time passes the system moves down to 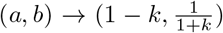. For *k* ≥ 1, trajectories move to 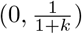. Note that this is the expected case, as the splicing of mRNA occurs in general faster than its degradation. Note that, in this case, trajectories cross below the green line, explaining why two solutions can be found for a single value of (*a, b*). The speed at which the system follows these trajectories depends on *γ*.

### 3.2 Real data

In order to assess the performance of the method on real data, we applied our method on the 4sU labeling experiment described in [24]. Briefly, mouse embryonic stem cells were plated at a density of 40000 cells/cm^2^ on gelatin-coated 10cm tissue culture plates and grown for approximately 14 hours. After addition of 4sU to the growth medium, cells were incubated at 37C for 10 minutes (10 minutes labeling pulse). RNA was then extracted and processed according to the protocol described in [25]. Reads that did not map to mouse ribosomal RNA sequences were aligned to intronic and exonic sequences using STAR V2.5 [26] and quantified using RSEM V1.1.17 [27], yielding intron and exon expression levels for unlabeled and labeled RNA.

For a single sample, the observable space represented in Fig 2 (center and right) is represented (in log coordinates) in Fig 3, left. We see that, while the points are centered on the expected region of the observable space, many transcripts lie below the diagonal or above the *r_l_* = 1 (or log(*r_l_*) = 0) line (in blue), which is not compatible with our model. We observe that those incompatible transcripts lying above the *r_l_* > 1 line are expressed at a much lower level than the transcripts lying below this line (see inset). A lower signal to noise ratio in low expressed genes could explain this difference. However, another likely explanation pertains to the fact that co-transcriptional processing is not accounted for by the Zeisel model. While it has been documented that an RNA molecule is often processed while being synthesized (the “assembly-line model”) [22], the Zeisel model considers synthesis and processing as two independent point events. This discrepancy is likely to be more relevant for short-lived (and thus low-expressed) transcripts, a sizeable fraction of which is expected to be nascent at sequencing time. Those nascent transcripts may contribute to an intron to exon ratios higher than one when they are incompletely synthesized (for example if the last exon has not yet been produced). This hypothesis is corroborated by considering unspliced transcripts length, which putatively affects synthesis time and thus the probability of being nascent at sequencing time. Transcripts lying above the *r_l_* > 1 line are indeed longer than those lying below this line (p – value < 10^−100^, Wilcoxon test).

**Figure 3:**
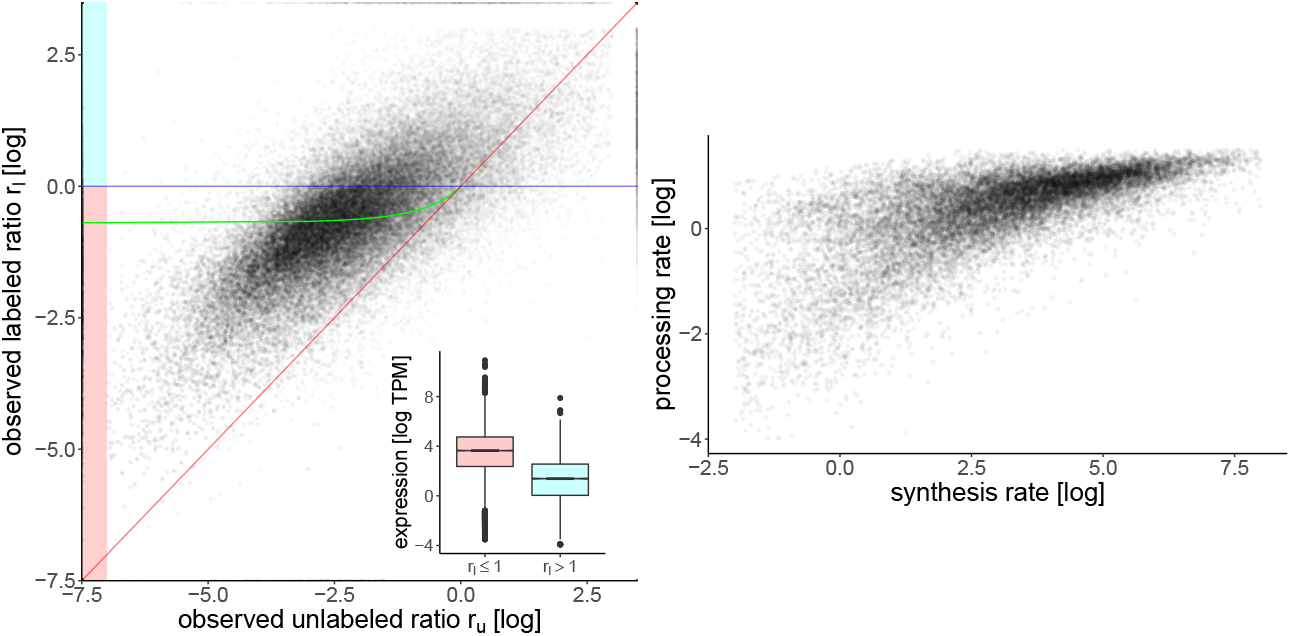
Real data. Left: Each point corresponds to a transcript with its transparency reflecting log expression value. Like in the previous figure, the green line is defined *y* = (2 – *x*)^−1^. For transcripts lying between the abscissa (in blue) and the green line, estimates of processing and degradation rates can be obtained by solving (12). For transcripts lying between the diagonal (in red) and the green line, estimates can be obtained by minimizing (15). The observed ratios for the remaining transcripts are not coherent with the model and are discarded. These trancripts (above the blue line) are lowly expressed compared to the ones below the blue line (see inset). Right:RNA processing rates are highly correlated to the synthesis rates (72%), which is consistent with co-transcriptional RNA processing.

The transcripts incompatible with our model, amount to 25% of protein-coding genes with an exon TPM higher than 1, and are discarded from further analyses. The processing and degradation rates were computed either by solving (12) when *r_l_* > (2 –*r_u_*)^−1^ or by optimizing (15) otherwise. For these cases that had two solutions (6% of the transcripts), we selected the one corresponding to rates most consistent with the other transcripts.

The resulting synthesis and processing rates are depicted in Fig.3, right. Although processing rates span a smaller range of values, they are highly correlated (72%), which is not surprising as RNA processing occurs co-transciptionaly [22]. More remarkable is the correlation of synthesis and degradation rates, displayed in Fig. 4, left. At 65%, it is very similar to the 66% reported by [21] for the same cell type. This is also consistent with the emerging concept of a coupling between RNA transcription and decay [28]. Our data indicate that genes span a large range of dynamics, irrespective of their expression level. Indeed, genes with high synthesis and degradation rates can have the same steady-state expression level as genes with low synthesis and degradation rates. However, the former will reach this steady state faster than the latter. It thus makes sense to consider our RNA metabolic rates in the functional frame of reference indicated in Fig. 4, left. One axis corresponds to the steady state RNA abundance, given by the log-ratio of synthesis over degradation rates (or equivalently by the difference of log of the rates). The second axis correspond to the responsiveness of the gene, i.e. how fast it reaches steady state (computed by the sum of the log of the synthesis and degradation rates). It has been observed before that genes involved in more reactive and dynamic biological processes such as chromatin remodeling or transcription regulation tend to have a higher turnover than genes involved in more stable processes such as basic metabolism [9]. We checked that our data confirm this observation by looking at the Gene Ontology (GO, [29])) annotations of biological processes most associated by [9] with high and low turnover, namely “transcription” and “monosaccharide metabolism”. Despite having similar steady-state abundances, transcripts of genes involved in transcription indeed have significantly faster dynamics and the ones involved in monosaccharide metabolism have significantly slower dynamics than the rest of the genes, as illustrated by the squares in Fig. 4, left and right. Other categories where our data confirms faster genes include chromatin modifications, cell cycle and transcription regulation.

**Figure 4:**
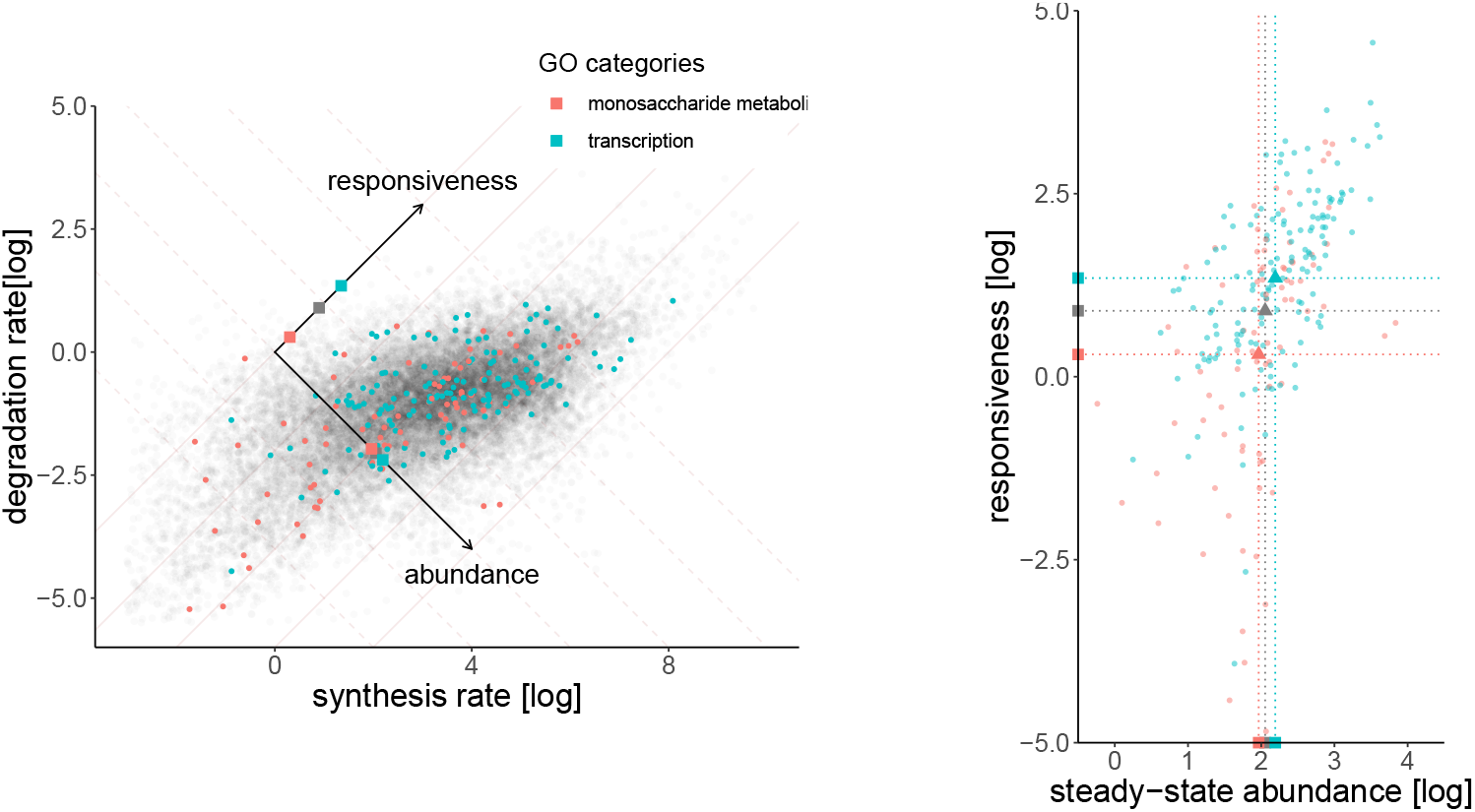
Left: Estimated RNA synthesis and degradation rates obtained from a single sample. These rates can also be considered in a different and maybe functionally more relevant frame of reference defined by the steady state abundance (first axis) and gene responsiveness (second and perpendicular axis), as illustrated by the background grid. Genes involved in fast adapting biological processes (such as transcription) tend to be more responsive than genes involved in stable functions (such as monosaccharide metabolism). The squares on the axes represent the projections of the mean rates for the respective categories (gray representing genes that belong to neither of the two categories) and indicate that mean transcript responsiveness (but not abundance) is strongly affected by the category. These two GO categories were selected for illustration because they were previously reported to be mostly enriched in high and low turn-over genes respectively [9]. Right: Same data as in left, but rotated and showing only colored dots, for visibility.

We assessed the precision of our method by comparing the resulting degradation rates to those published for the same cell type by [21]. Those were obtained by using three replicates and seven time points and applying the SLAM-seq nucleotide-conversion method that, unlike metabolic labeling, does not require biochemical separation between the labeled and unlabeled RNA and is thus not affected by noise generated by the imperfect separation process (although that method has its own source of noise). From our data, we obtained gene degradation rates by taking, for each gene, the weighted average degradation rates of the corresponding transcripts The weights were given by the mean exonic expression levels (unlabeled and labeled). We expect a lower precision for transcripts close to the *r_l_* = 1 line, for which the labeling time was likely somewhat too short, so to assess the correlation, we weighted the transcripts by 1 – *r_l_*. Fig. 5, left, compares degradation rates obtained in our experiments with those reported by [21], keeping only genes with an average expression value higher than 100 TPM. We expect a higher precision for highly expressed genes, as this allows for a more precises estimates of the intron to exon ratios. This is indeed the case, and depending on the expression threshold and the sample, the correlation between our data and the previously published rates, we obtain a correlation ranging between 30% and 67% for a single sample estimate (see Fig. 5, left). As these experiments were performed in different labs using different methods, these numbers show that our rates obtained on a single sample and time point are meaningful. For comparison, [30] report correlations around 70% by using the *same* data, but changing only the method of analysis. Using three replicates, [25] report a 26% correlation using the INSPEcT package.

**Figure 5:**
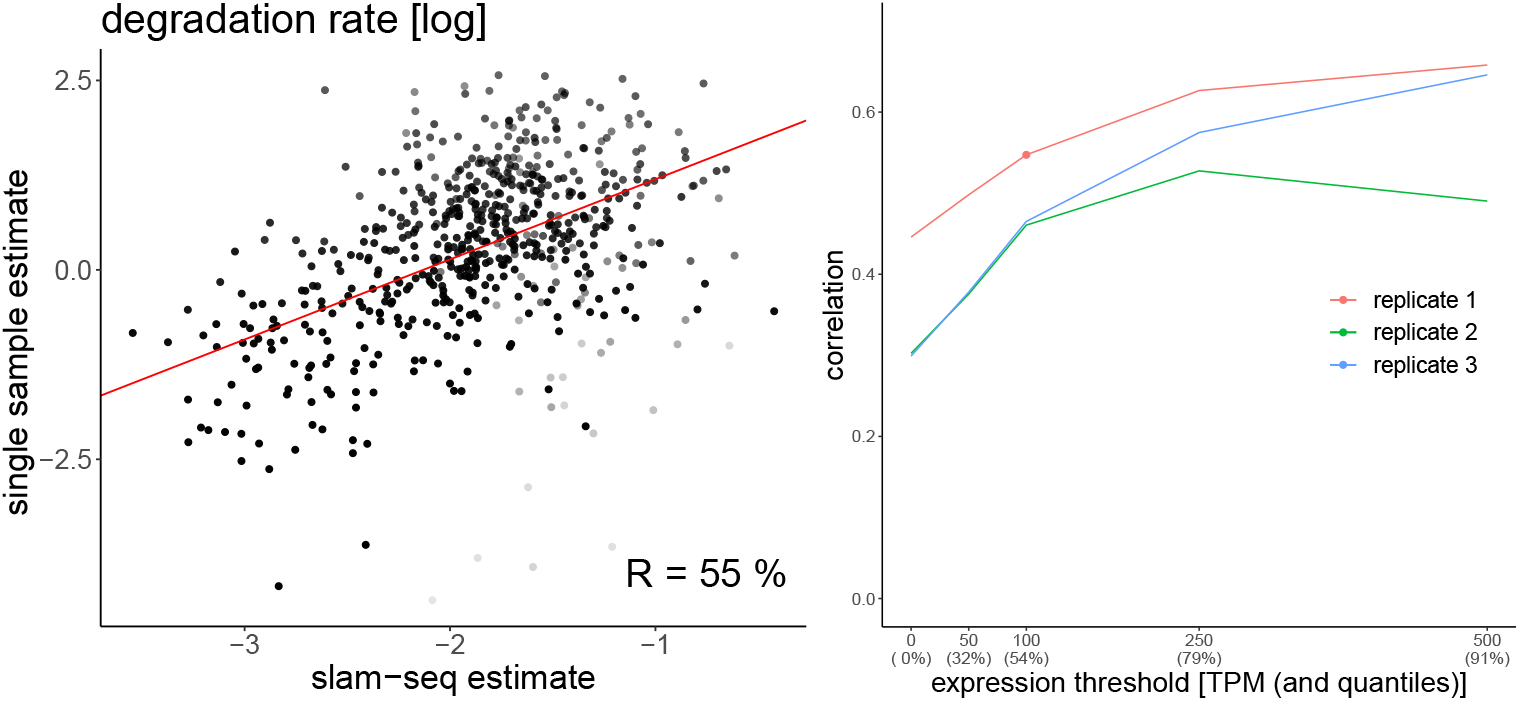
Left: degradation rates estimated from a single sample plotted against degradation rates published in [21] (obtained using slam-seq). The red line is obtained through weighted linear regression. The weights are set as 1 –*r_l_* as indicated by the transparency of the dots. The (weighted) correlation of 55% indicates that the estimated rates are meaningful. Only genes with a mean exon TPM above 100 are taken into account. Right: Correlation between degradation rates obtained by [21] and the ones obtained our single-sample method as a function of expression level. Each line represents a biological replicate. The dot corresponds to the data shown on the left. As expected, the correlation is higher for highly expressed genes, as the intro to exon ratios can be more reliably estimated. In this experiment, replicate 1 correlates better than the two others, indicating that it is probably of better quality.

### 3.3 Comparison with INSPEcT

Since our method estimates metabolic rates from a single sample, we decided to compare its results to the “initial guess” provided by the INSPEcT method, to our knowledge the only other method that does not need multiple samples. Note however, that those rates are only the initial step of the INSPEcT method, and should not be confused with the global outcome of INSPEcT, which then aggregates multiple samples for the estimation. For concision, we will in the section refer to our method as SSRE (Single Sample Rate Estimation). The main differences between the two approaches is that INSPEcT assumes no degradation on labeled RNA and requires the estimation of a scaling factor accounting for the difference in RNA concentration between labeled and unlabeld samples, which SSRE avoids by considering the intron to exon ratio in each sample separately for each sample. Furthermore, INSPEcT requires the estimation the time derivative of the RNA abundances, which is avoided in SSRE by taking advantage of the analytical solution to the Zeisel model.

We used the INSPEcT package for R (more specifically the `newINSPEcT` function with parameter `preexisting=TRUE` and the `ratesFirstGuess` function) on the same data. The expression variance required by this function was estimated from the expression level from all three replicates using Loess regression. It took about 10 minutes to estimates rates for each replicate (about 50000 transcripts) using a single 2.8GHz core from a laptop computer, whereas our implementation of SSRE took about 30 seconds to complete on the same machine.

In addition to direct rate comparison, we decided to compare the methods using three criteria: (1) correlation with published rate, (2) rate distributions and (3) reproducibility across replicates. Figure 6 shows that the two methods provides synthesis and processing rates that are well correlated, while degradation rates are not. Moreover, the degradation rates obtained by INSPEcT correlate well to previously published rates only for one of the three samples. In contrast to degradation rates obtained with SSRE, the correlation with previously publised rates does not improve when focusing on highly expressed genes, it even become negative for replicates 1 and 3 (data not shown). This suggests that for SSRE, degradation rate estimation is likely to improve with higher sequencing depth (and thus a more precise estimate of the intron to exon ratio). Finally, the rates computed using the INSPEcT method do not exhibit the previously documented correlation between synthesis and degradation rates [21]. This leads us to think that our degradation rates are closer to the real rates than the ones provided by the INSPEcT “first guess”. This should not come as a surprised, as our method does not assume that labeled RNA does not degrade and estimates degradation and processing rates independently from the synthesis rate.

**Figure 6:**
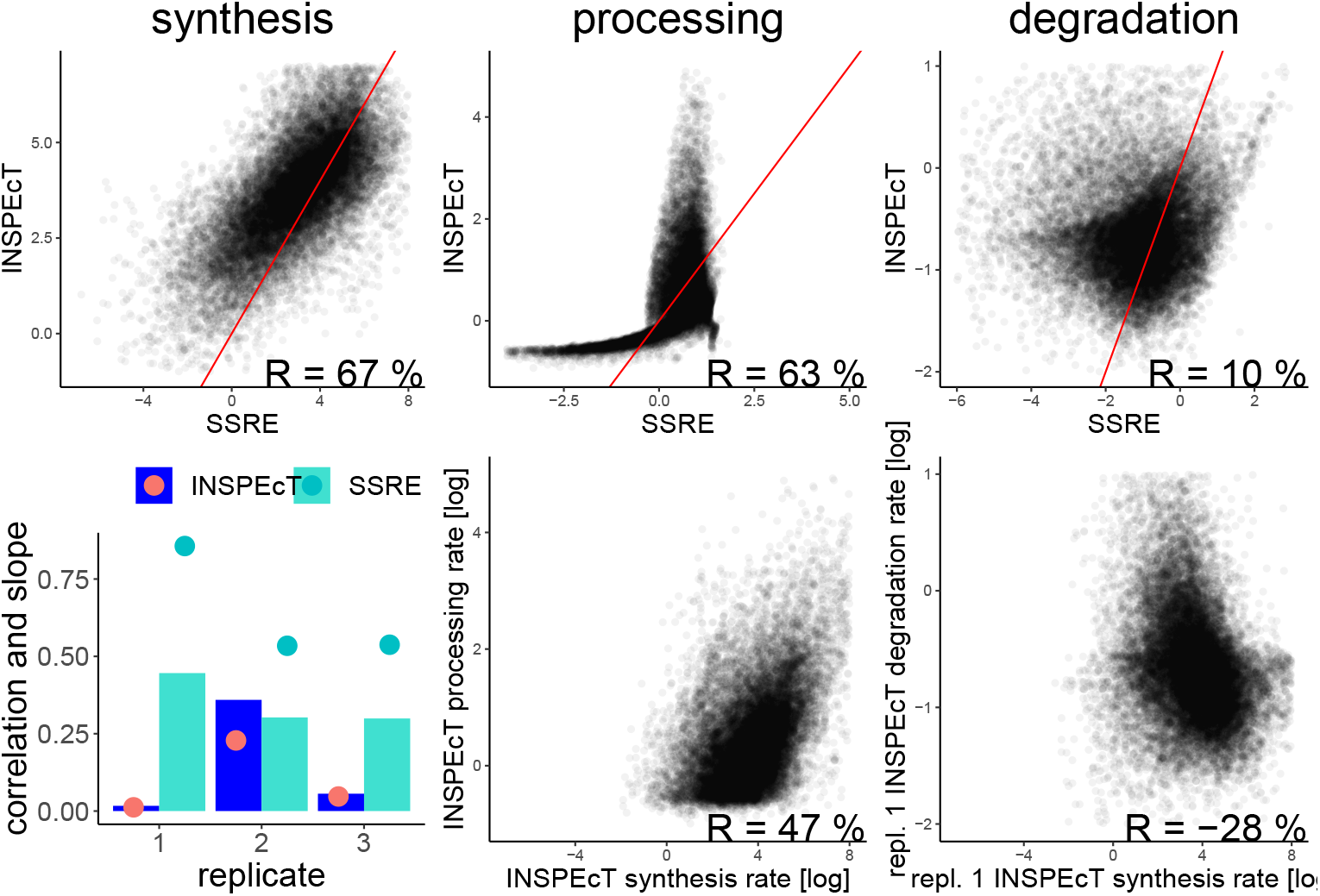
Comparion of our method (SSRE) with the INSPEcT “first guess” on th same data. Top row: direct comparison of rates obtained with our method and with the INSPEcT package on a single sample. Synthesis and processing rates are well correlated but not the degradation rate (Spearman correlation shown). The red bar indicates the diagonal. Bottom left: bars indicate the correlation of degradation rates with previously published data [21], as in Fig. 5. The INSPEcT method provides degradation rates with good correlation only for one of the three replicates (repl. 2), whereas it is the case for all three replicates using our method. The dots indicate the slope of the regression line in log-log space (as in Fig. 5, left). Slopes obtained from SSRE estimates are closer to one, which correspond to the ideal case of a linear relation ship between the (non-log) rates. Bottom center and right: Rates obtained with INSPEcT also reproduce the positive correlation between synthesis and processing rates, but they produce a negative correlation between synthesis and degradation rates, unlike our method and previously published results.

The synthesis and processing rates provided by the two methods are relatively well correlated, and INSPEcT provides rates that are more consistants across replicates (see Supplementary Fig D.8.) It is also interesting to note that SSRE tends to show an upper bound for the processing rate, while INSPEcT first shows a lower bound for that rate. It is difficult to speculate which (if any) is more likely true, but an upper bound would be consistent with biophysical constraints in a leaky co-transcriptional RNA splicing setting. Fig D.8 also shows that unlike INSPEcT, SSRE computes degradation rates that span a larger range of values than processing rates, a property also reported in [11] for a different system.

## 4 Discussion

In this paper, we presented a method to estimate synthesis, processing and degradation rates of RNA transcripts from a single 4sU labeled sample. We validated our method first *in silico* and then on real data obtained from mouse embryonic stem cells. Using our method we first replicated, on a different cell type, previous findings about the enrichment in high or low turn-over genes of specific cellular processes. Second, we showed that the rates obtained with our method correlate well (between 30% and 67%) with published rates obtained by applying SLAM-seq to the same cell types. Methods for such estimation have been published before, but they usually require a sufficient number of samples (around a dozen). We compared our method to the initial step of the INSPEcT method, which handles each sample separately, and obtain similar synthesis and processing rates, but different degradation rates. Our rates correlate more consistently with previously published degradation rates obtained with nuclear conversion methods on the same system, and even more so for highly expressed transcripts. Rates obtain with our method also better reproduce previously observed statistical relationships between rates, although synthesis and processing rates are less consistant across replicates. Taken together these results suggest that our method provides more reliable degradation rates.

In contrast to other methods, our method explicitly uses the analytical solution to the Zeisel model of RNA dynamics. Moreover, our method is self-normalizing as it only uses the ratio of intron to exon expression levels. It is thus not affected by differences in sequencing depth of the various samples. This approach makes our method also faster than other methods as it boils down to numerically solving on a bounded domain either a univariate equation or a one-dimensional optimization for each transcript. Our method could thus be a suitable alternative to the initial step of the INSPEcT method especially when using a large number of samples as it is also about 20 times faster.

Similarly to the initial step of the INSPEcT method, a caveat of our method is that a sizable fraction of mostly lowly expressed transcripts (about 25 % in our case) are inconsistent with the model and their dynamics cannot be estimated. Together with the high correlation between synthesis and processing rate, it suggests that modeling transcription and processing as independent events is a simplification that could be reconsidered, as the coupling between the two has been documented [22]. However, this limitation of the Zeisel model is likely to also affect other methods using it [20, 31].

Another limitation of the method is that, unlike in [20], it does not consider the effect of leakage of unlabeled RNA in the labeled RNA pool because of unspecific capture. This leakage has the effect of reducing *r_l_* towards the diagonal, and could potentially be estimated from the data as it is shared across all transcripts. Another improvement would be to embed this method in a probabilistic framework in order to quantify the estimate uncertainty (as in [30] for a simpler model) or to determine the optimal labeling time (as in [32]).

While using a single sample allows to reduce costs, this is not the only merit of this approach. In practice most experiments will have biological replicates, in which case our methods enables obtaining point estimates of *α, β* and *γ* for each of them. This in turn allows for estimating their variance, as well as assessing sample quality (e.g. if one of them systematically gives very different estimates for all genes). Moreover, because cell growth is likely to be limited during (short) labeling time, it is less likely to interfere in the estimation process than when using time course data, where it can have an effet [18]. In addition, when used in a time-course experiment, our method allows to investigate the evolution of these rates over time and assess whether these rates are stationnary. Finally, the theoretical results obtained in this paper, could be used to improve other methods. For example, the method could be used to analyze SLAM-seq data which would reduce the number of samples but also provide estimate for the processing rate. Another possible application is single cell RNA velocity [31], where the Zeisel model of RNA dynamics is also used, but splicing rates 7 are set to be equal for all transcripts. While it has been documented (and is consistent with our data) that splicing rates are more homogeneous than degradation rates [11], this is potentially an approximation that could be improved with our framework to increase the accuracy of the method, for example by considering the strong correlation between the synthesis and processing rates. Finally, our method could also be used in conjunction with the recent developements in single cell metabolic labeling experiments [33, 34].

## Acknowledgements

This work was funded by the Swiss National Science Foundation through grant no. FN 310030_152724/1 to S.B and PP00P3_150667 and the NCCR in RNA & Disease to A.C.M.

## Availability of data and code

An R package implementing our method is available on github, together with the code used to generate the figures as well as the gene expression data used: https://github.com/BergmannLab/SingleSampleRNAdynamics

The raw data files data are available on the Gene Expression Omnibus accession number GEO:GSE150286 (main replicate) and GEO: GSE143277 (second and third replicates of Fig. 5, right and Supplementary Fig. D.8.

## Author’s contribution

M.H. developed and implemented the method, analyzed the expression quantification data, interpreted the results, figured out the proof, generated the figures and wrote the manuscript, A.B. performed the experiments and interpreted the results, A.C.M. initiated the project, designed the study and generated the expression quantification data, S.B. reviewed the math, interpreted the results and supervised the process. All authors contributed to the manuscript.

## A Derivation of the model solution

This is a first order linear ordinary differential equation in p(t) and m(t) that can be expressed in matrix form as

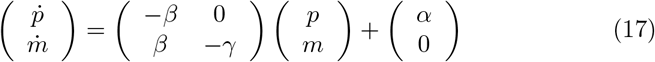

The solution to this equation is given

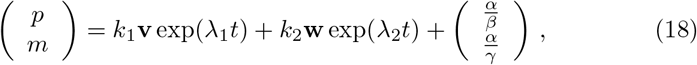

by where *k*_1_ and *k*_2_ are scalar constants determined by the boundary conditions, λ_1_, λ_2_ are eigenvalues of the matrix in (17) and **v**, **w** are the corresponding eigenvectors.

The eigenvalues are given by λ_1_ = –*β* and λ_2_ = –*γ*. The first eigenvector **v** is obtained by solving

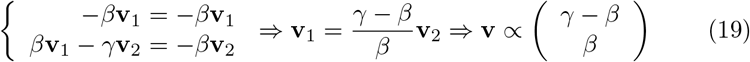

Similarly the second eigenvector is obtained by solving

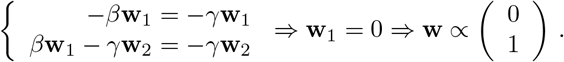

The solution to (17) is thus given by

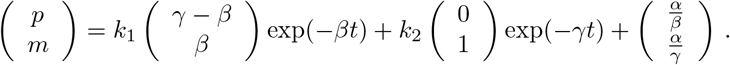

Expressed by its component this is equivalent to

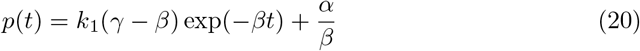

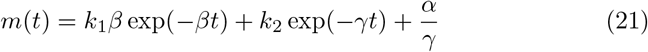

We now turn to the boundary conditions to determine *k*_1_ and *k*_2_. The boundary conditions are different for the unlabeled and the labeled RNA.

### Unlabeled RNA

Like in [19], we assume the system to be in steady-state prior to labeling. The steady-state is given by solving (17) with 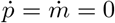.

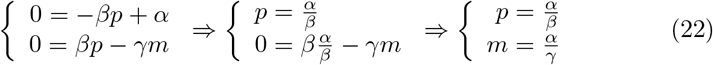

During labeling time, we assume that no unlabeled RNA is synthesized such that *α* = 0. Assuming that we start labeling at time *t* = 0, we thus have

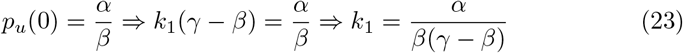

Moreover we have

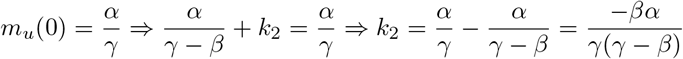

This leads us to the solution for the unlabeled RNA

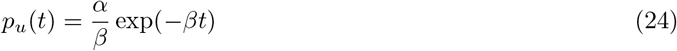

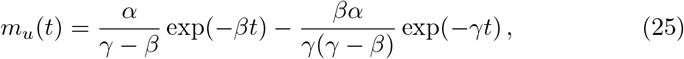

where the *u* label indicates that this corresponds to the unlabeled RNA pool.

### Labeled RNA

The solution for the labeled RNA could be obtained the same way as for the unlabeled RNA, but setting *α* ≠ 0 and *p_l_* (0) = *m_l_*(0) = 0. However, it is simpler to notice that the total RNA (labeled and non-labeled) stay at steady-state during the labeling such that we have the following solution for labeled RNA.

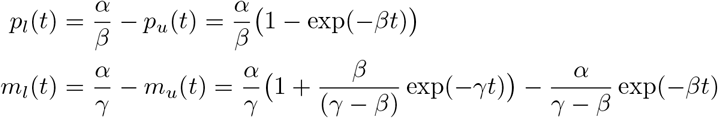

where the *l* label indicates that this corresponds to the labeled RNA pool.

## B Equation simplification

In this appendix we show how the the system given by equations (10, 11) can be simplified to Eq. (12) to infer the ratio *k* between the processing and degradation rate. Starting from

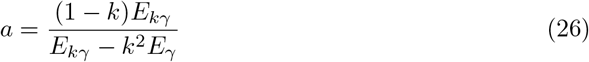

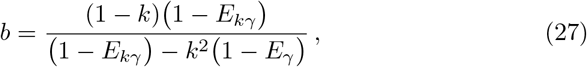

we have

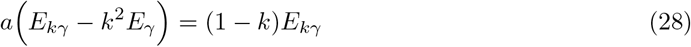

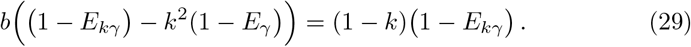

Summing (28) and (29) yields

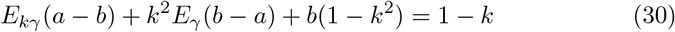

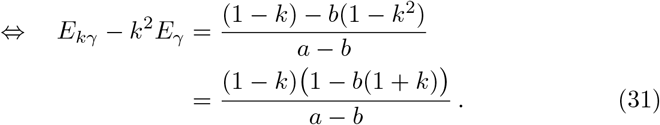

Dividing (28) by (29) and inserting (31) results in

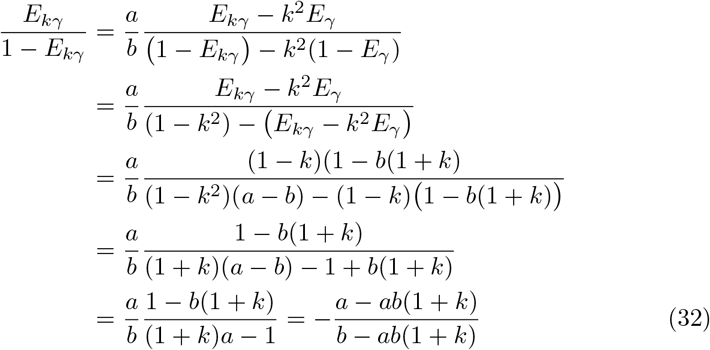

It follows that

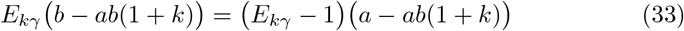

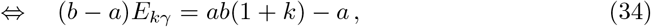

an thus

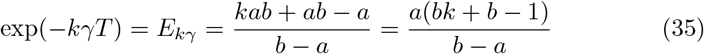

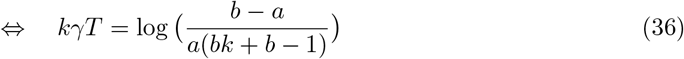

Moreover, from (10), we have that

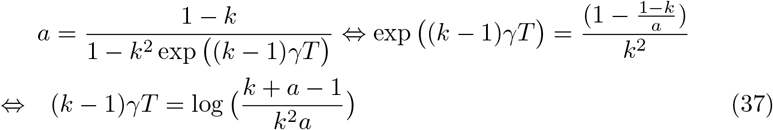

Multiplying (36) by 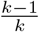 and subtracting (37) results in

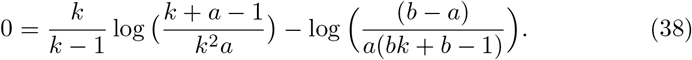

This equation also provides upper and lower bounds for *k* as both 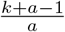 and *bk* + *b* – 1 must be strictly positive for their logarithm to be defined and

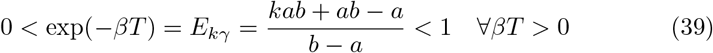

for (35) to hold. Developing these three conditions results in the following domain of definition 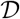 for *k*:

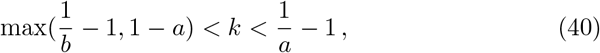

where 0 < *a* < *b* < 1.

## C Proof of unicity of solution

In this appendix, we prove that (12) has a single solution for *b* > (2 – *a*)^−1^. We first note that 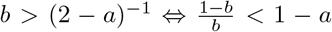, so the lower bound for *k* is *k*___ − = 1 – *a*. We then define the right-hand side of (12) as

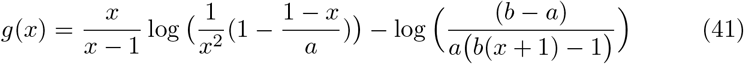

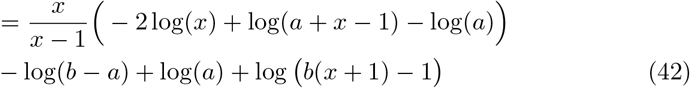

We then observe that on that lower bound 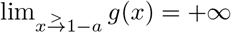 because *a* + *x* − 1 tends to zero and *x* – 1 is negative.

On the other hand, for the upper bound 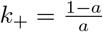, we have

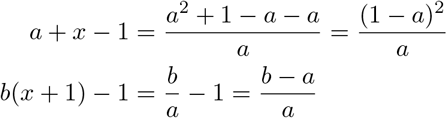

we can deduce that the upper bound of *x* is a zero of *g*:

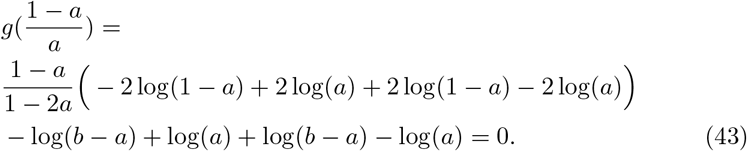

Moreover, the derivative of *g* is given by:

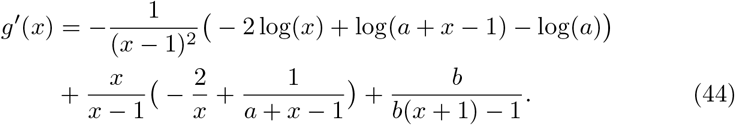

Then

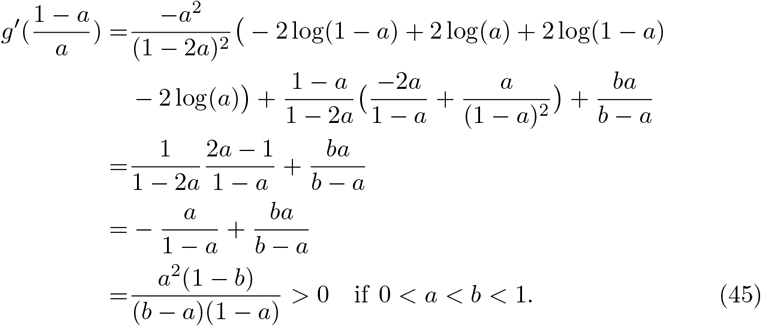

Since *g*(*k*_+_) reaches zero from below, while *g*(*k*_−_) > 0, we can infer that *g*(*x*) has a zero between *k*_−_ and *k*_+_ as illustrated on Fig C.7.

To show that this zero is unique, we look at the sign of *g*′(*x*).

We can rewrite *g*′(*x*) as

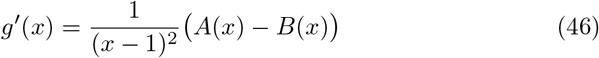

where

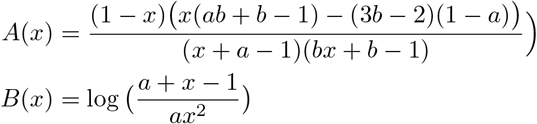

Let *x*_0_ be a zero of *g*′, i.e., the position of a local extrema of *g*. We have

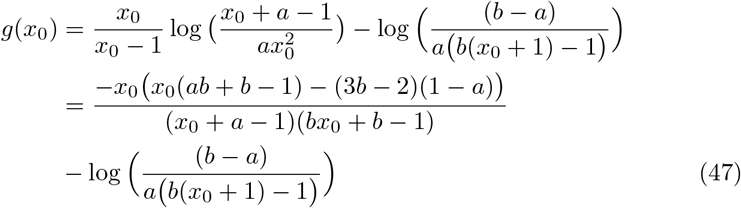

The second equality holds because *g*′(*x*_0_) = 0 by definition of *x*_0_. By multiplying (47) by (*bx*_0_ + *b* – 1), which is positive, we can then define a new function *h*(*x*) whose sign is the same as the sign of *g*(*x*) for *x* = *x*_0_ (see Fig C.7 for an illustration).

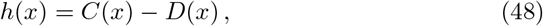

where

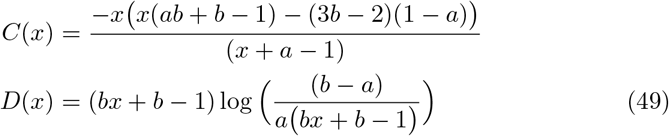

**Figure C.7:**
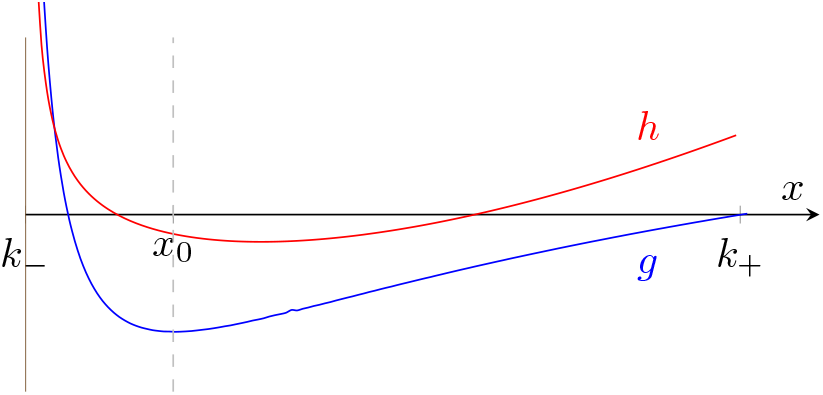
Sketch of the proof that *g*(*x*) has a single zero in 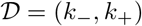. We first show that lim_*x*→*k*____*g*(*x*) = ∞, that lim_*x*→*k*_+__*g*(*x*) = 0 and lim_*x*→*k*_+__*g*′(*x*) > 0, so that *g* must cross the x-axis on 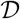. To show that it only does it once, we consider a function *h*(*x*) that has the same sign as *g*(*x*) when *g*′(*x*) = 0. We show that *h* is convex on 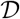 and thus *g* cannot have a negative extrema, followed by a positive extrema, followed by a negative extrema. Hence it cannot have more than one zero on 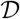.

We can now compute the second derivatives of *C*(*x*) and *D*(*x*).

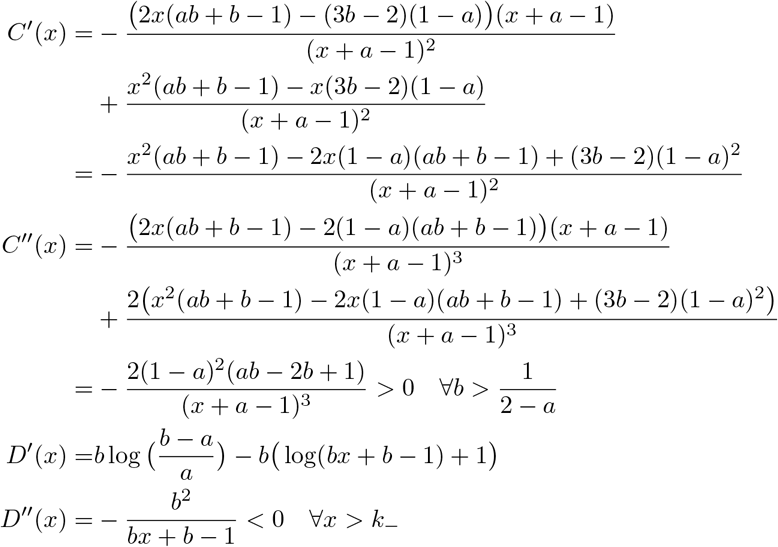

Hence

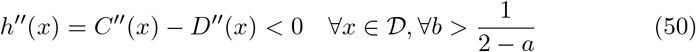

This means that *h* is convex, so there cannot be three points *x*_1_ < *x*_2_ < *x*_3_ such that 0 > *h*(*x*_1_) < *h*(*x*_2_) > 0 > *h*(*x*_3_). Hence the same can be said of three zeros of *g*′, so *g*(*x*) cannot have more that one zero. □

## D Supplementary figure

**Figure D.8:**
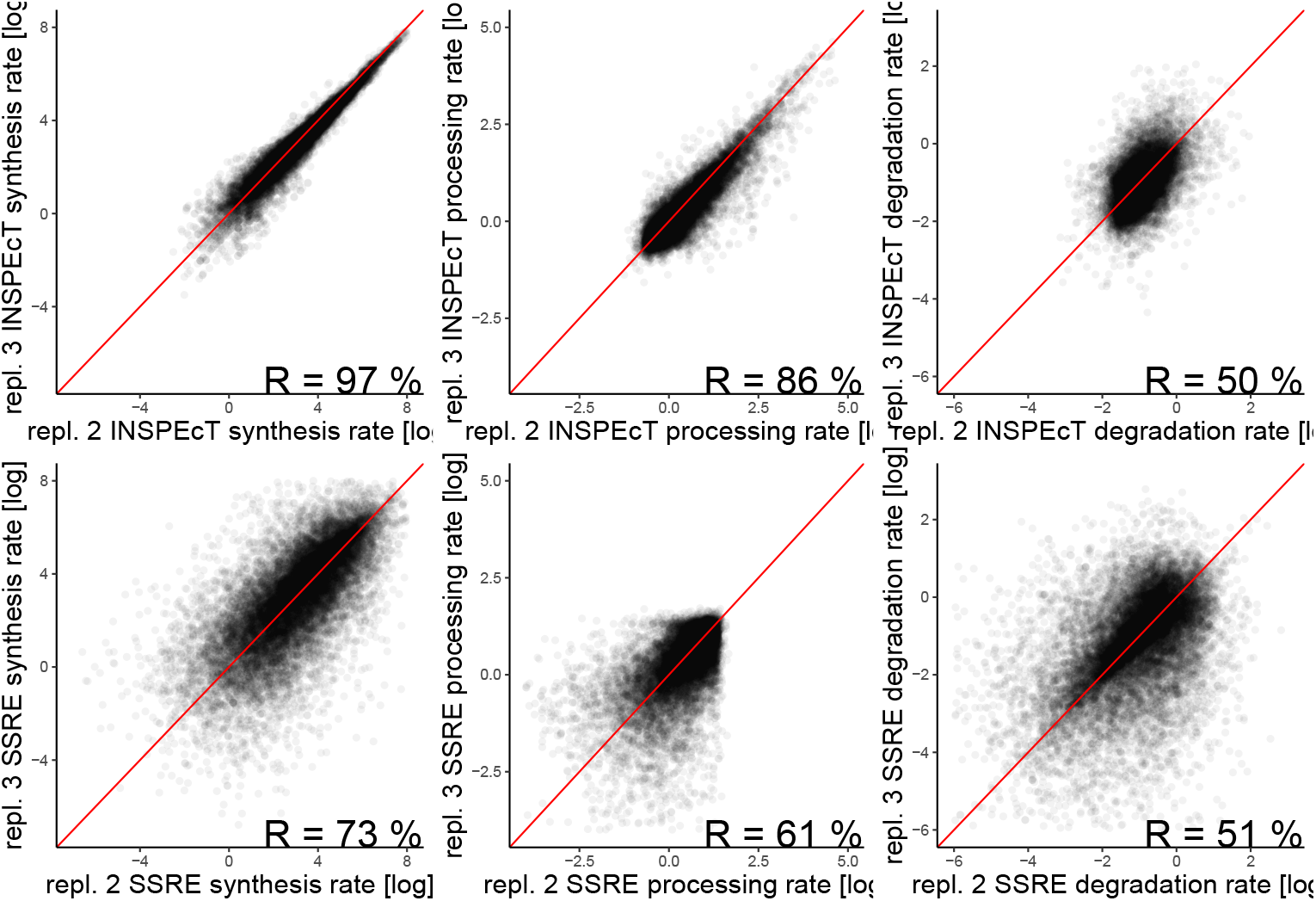
Correlation of rates across two replicates. Top row: Rates obtained with INSPEcT. Bottom row: rates obtained with our method. The INSPEcT method provides rates that are more consistent across replicates for synthesis and processing rates, but not for degradation rates. Spearman correlation is indicated and the red line shows the diagonal.

